# High-Throughput Profiling of Bacterial Respiration Using the Resipher Reveals Functional Responses to Nutrients and Antibiotics

**DOI:** 10.64898/2026.06.17.732880

**Authors:** Nimra Khalid, Aria Eshraghi

**Affiliations:** Department of Infectious Diseases & Immunology, University of Florida, Gainesville, Florida, USA; Emerging Pathogens Institute, University of Florida, Gainesville, Florida; Department of Oral Biology, University of Florida, Gainesville, Florida

## Abstract

Oxygen consumption is a direct functional readout of bacterial respiration and metabolic state, yet existing methods for quantifying oxygen dynamics are limited in throughput and temporal resolution. Here, we establish a high-throughput platform for real-time profiling of bacterial respiration by adapting the Resipher, a non-invasive oxygen quantification system, for use in bacterial cultures. Measurements obtained with the Resipher were comparable to those generated using a Clark-type electrode-based high-resolution respirometer, validating its quantitative accuracy. Across Gram-negative (*Escherichia coli, Francisella novicida*) and Gram-positive (*Enterococcus faecalis, Staphylococcus aureus*) species, the Resipher generated reproducible measurements under both growth-permissive and growth-limited conditions, enabling assessment of respiration, independent of proliferation. Functional profiling revealed that oxygen consumption responds dynamically to nutrient availability and electron transport chain perturbation, including species-specific inhibition by benzarone. Notably, oxygen consumption profiles distinguished bactericidal and bacteriostatic antibiotics, with bactericidal agents transiently increasing respiration and bacteriostatic agents suppressing metabolic activity. Together, these findings establish oxygen consumption as a sensitive physiological readout and highlight the potential utility of respiratory profiling for mechanistic studies.

## INTRODUCTION

Oxygen plays a fundamental role in physiology, metabolism, and pathogenesis in bacteria^1,2^. As a terminal electron acceptor in aerobic respiration, oxygen availability directly impacts energy generation, redox balance, and regulation of metabolic pathways^3,4^. Shifts in oxygen levels can also trigger profound changes in bacterial enzyme activity, and virulence gene expression, thereby influencing host colonization, survival, and pathogenesis^5–7^. Likewise, varying environmental and genetic conditions lead to dynamic changes in respiration^8–12^. Thus,quantifying oxygen consumption rates (OCR) in bacteria is a powerful tool to investigate fundamental physiological processes.

Aerobic respiration (hereafter referred to as respiration) is the most efficient form of energy production, generating up to 38 ATP molecules from the oxidation of one glucose molecule^13^. In bacterial respiration, the electron transport chain (ETC) facilitates transfer of electrons to a terminal electron acceptor, resulting in translocation of protons across the cytoplasmic membrane into the periplasm^14^. ATP synthase uses this chemiosmotic gradient to phosphorylate ADP to generate ATP. While facultative anaerobes can utilize alternative electron acceptors when oxygen is limited, oxygen is the only terminal electron acceptor in aerobic bacteria^15^. Therefore, oxygen consumption is directly linked to ATP synthesis in aerobic respiration. Consequently, a decline in dissolved oxygen in bacterial cultures serves as a reliable measure of respiratory activity^16–18^. Accurate and robust techniques for monitoring oxygen consumption are required to advance our knowledge of microbial physiology and pathogenesis.

Dynamic measurement of oxygen utilization can provide insights into bacterial energy demands under stress-induced conditions. For example, antimicrobials have profound effects on bacterial oxygen consumption, indicating alterations to ETC activity and shifts in metabolism^19^. These effects can be disparate. Attenuated oxygen utilization following tetracycline is linked with metabolic suppression^12^. Conversely, increased oxygen consumption after exposure to bactericidal drugs may result from adaptive stress responses^12,20,21^. Similarly, bacterial metabolic pathways shift in response to nutrient scarcity, resulting in reduced ETC activity and decreased OCR^8^. Genetic changes can also modulate bacterial oxygen consumption and induce changes in redox balance^22,23^. Therefore, monitoring oxygen consumption is a powerful tool for characterizing bacterial metabolic status under different environmental and genetic conditions.

Several techniques have been developed to quantify oxygen consumption in living cells; however, current technologies lack temporal resolution, are invasive, or require extensive sample manipulation, limiting their utility in high-throughput time course studies (Table S1). Many of these methods are limited to the analysis of single samples and are not suitable for continuous oxygen monitoring^16^. Traditional polarographic sensors, such as the Clark-type electrodes, pose challenges in accurate measurement in small-volumes because of direct oxygen consumption by the electrode. These sensors also need constant shaking during the reading and require repeated recalibration to avoid electrode drift^17^. Similarly, chemical assays to quantify dissolved oxygen, such the Winkler method, require multiple labor-intensive steps and careful titration that make rapid assays unfeasible^24–26^. While more advanced equipment, such as the Oroboros O2k, provide precise measurements of oxygen consumption, their low-throughput and labor-intensive protocols limit their utility for monitoring metabolic dynamics^27^. In contrast, the Seahorse platform (Agilent Technologies) offers high-throughput measurement of oxygen consumption. However, both the equipment and consumables have high costs and running the instrument requires significant technical expertise. Furthermore, the limitations on the duration of assays pose challenges for long-term studies^28^. This highlights the demand for innovative tools that provide precise, continuous, and high-throughput measurements of cellular oxygen consumption.

The Resipher is a non-invasive oxygen quantification system that is designed to provide high-throughput concentration data from 96-well plates. The equipment is comprised of a sensor that attaches to a sterile lid with oxygen-sensitive tips that extend into each well of a 96-well plate. The sensor excites and records emission from a proprietary oxygen-sensitive dye on the tips which are submerged in the media. This dye emits light with a characteristic lifetime after excitation, and molecular oxygen shortens this lifetime by collisional quenching. By measuring the luminescence decay time, oxygen concentration can be determined reliably and independent of signal intensity. Measurements can be performed in rapid succession to provide continuous monitoring of oxygen concentration that is reported to a computer in real-time. The Resipher has been used extensively to measure oxygen concentration in eukaryotes, such as yeasts, mammalian cells and invertebrates like *Caenorhabditis elegans*; however, its use in microbiology research is limited^29–32^. Measuring oxygen concentration in fast-growing bacterial cultures is a challenging task because their relatively short doubling time requires rapid measurements in short succession. High-throughput measurements are even more challenging because the small volumes of samples result in short times to nutrient limitations and skewed metabolic data^33^. An advantage of the Resipher is that the plates are not sealed, allowing oxygen to readily diffuse into the media and allowing for both long and short-term studies. While the primary use of the Resipher has been on eukaryotes, such as mammalian cell culture studies, recent interest in microbial metabolism has prompted a need to extend its applications to bacteria, though this area is still being explored and optimized. The combination of continuous monitoring, high-throughput capacity, precision, and user-friendly operation make the Resipher a powerful tool in microbial metabolic research.

In this study, we present a detailed assessment of the Resipher for measuring oxygen consumption in diverse bacterial species. Although this work establishes an experimental framework for quantifying bacterial respiration using the Resipher, our goal extends beyond technical validation. Respiratory activity is a central determinant of bacterial physiology, linking nutrient utilization, energy generation, stress adaptation, and antimicrobial susceptibility. Consequently, the ability to monitor oxygen consumption with high temporal resolution and experimental throughput provides an opportunity to interrogate fundamental microbiological processes that are otherwise difficult to assess in a scalable manner. We therefore sought not only to evaluate the suitability of the Resipher for bacterial systems, but also to determine whether respiratory profiling can reveal biologically meaningful responses to environmental and pharmacological perturbations across diverse bacterial species. We demonstrate that the Resipher provides precise and reproducible measurements of bacterial oxygen consumption and, by optimizing culture conditions, we extend its use to evaluate metabolic responses to nutrient availability, electron transport chain perturbation, and antibiotic exposure. Our results support the utility of the Resipher as a versatile platform for microbial metabolic profiling and highlight the broader value of respiratory phenotyping for studying bacterial physiology, antimicrobial mechanisms of action, and metabolic engineering.

## METHODS

### Bacterial strains and culture conditions

*Francisella novicida* Utah 112 was obtained from the NIH Biodefense and Emerging Infections Research Resources Repository (BEI resources, NR-13), *E. coli* K-12 MG1655 was obtained from Subhashinie Kariyawasam (University of Florida), and *Enterococcus faecalis* OG1RF and *Staphylococcus aureus* Newman were obtained from Jose Lemos (University of Florida). All bacterial strains were grown on tryptic soy agar (TSA) plates except for *F. novicida* for which TSA was supplemented with 0.1% cysteine (TSAC). For broth cultures, bacteria were grown in tryptic soy broth with or without 0.1% cysteine (TSBC/TSB), M9 minimal medium, and chemically defined media (CDM), as indicated^34,35^.

### Preparation of bacteria for oxygen quantification

To quantify oxygen in cultures during logarithmic growth, strains were grown overnight in TSB or TSBC for 16 ± 2 hours, diluted 1:100 to 1:500 in their respective media, grown to logarithmic phase (OD_600nm_ = 0.3 - 0.7), then cultures were normalized to OD_600nm_ = 0.01. To measure oxygen in conditions not permissive for bacterial growth, strains were streaked onto TSA/TSAC plates and grown for 14 ± 0.5 hours, scraped into media, washed, and resuspended in their respective media. OD_600nm_ was measured and adjusted to the concentrations indicated in the results (0.06 - 0.5). For carbon source supplementation experiments, bacteria were resuspended in basal media containing the indicated carbon sources. In these experiments, TSB-resuspended bacteria were included as control conditions to benchmark robust respiratory activity.

To assess the effects of ETC inhibitors and antibiotics, bacterial suspensions were exposed to the compounds immediately prior to quantifying oxygen. Stocks of the ETC inhibitors CCCP and benzarone were prepared in 100% DMSO at 20 mM and 10 mM, respectively. These were diluted to 25 µM CCCP (final DMSO concentration 0.1%) and 25 µM and 50 µM benzarone (final DMSO concentration 0.5%) in bacteriological media. Untreated bacteria were exposed to an identical concentration of DMSO to control for solvent-induced changes. Stock solutions of antibiotics were prepared as follows: 50 mg/mL enrofloxacin in 100% DMSO, 50 mg/mL ampicillin in water, 10 mg/mL doxycycline in 100% DMSO, 20mg/mL chloramphenicol in 50% ethanol, and 7.5 mg/mL tetracycline in 50% ethanol. For *E. coli*, treatments included enrofloxacin at one half and ten times the IC_90_ (0.15 µg/mL and 3 µg/mL, respectively) and ampicillin at one eighth the IC_90_ (18.8 µg/mL). *E. faecalis* was exposed to enrofloxacin at one half and five times the IC_90_ (4 µg/mL and 40 µg/mL, respectively), and to ampicillin at one eighth the IC_90_ (3.1 µg/mL). Bacteriostatic antibiotics were evaluated at their IC_90_ concentrations in both bacterial strains. *E. coli* was treated with doxycycline, chloramphenicol, and tetracycline at 5 µg/mL, 3 µg/mL, and 6 µg/mL, respectively, and for *E. faecalis*, the corresponding concentrations were 0.2 µg/mL, 6 µg/mL, and 0.5 µg/mL. Untreated bacteria were exposed to identical concentrations of solvent to control for solvent-induced changes. Unused outer wells were filled with media lacking bacteria to monitor environmental contamination. 100 µL of each condition was added to 96-well plates and oxygen concentration was measured with the Resipher.

### Oxygen quantification with the Resipher

The Resipher equipment was set up as recommended by the manufacturer (Lucid Scientific) to monitor oxygen levels in standard flat-bottom 96-well plates (Greiner Bio-One). Sensing lids that contain the oxygen probes were placed on top of the 96-well plates, such that the probes were submerged in the bacterial suspensions. These lids were attached to the corresponding sensors, and these assemblies were placed in a humidified incubator at either 37 °C or 25 °C without shaking. Probes were set to fixed height without oscillation to allow for total oxygen concentration measurements. The Resipher workstation quantified oxygen concentration, temperature, relative humidity, and atmospheric pressure. Oxygen consumption rates (µM/min) were determined using the following calculation: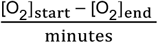 where [O_2_]_start_ is the baseline oxygen concentration and [O_2_]_end_ is the concentration measured after a duration of time (X minutes).

### Comparison of oxygen quantification with the Resipher and Oroboros O2k

Oxygen consumption rates were independently measured using an Oroboros O2k high-resolution respirometer equipped with a Clark-type polarographic oxygen sensor (OroboPOS). Logarithmic-phase bacterial cultures were normalized to OD_600nm_ = 0.2 and introduced into the respirometry chamber under conditions matched as closely as possible to those used for Resipher measurements. Oxygen consumption rates were calculated according to manufacturer guidelines and normalized to colony-forming units (µM/min/CFU) using the following calculation: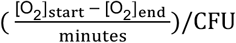

### Enumeration of colony-forming units

Colony-forming unit (CFU) enumeration was performed on the bacterial suspensions using standard methods. Briefly, bacterial suspensions at the start and end of the oxygen quantification were serially diluted in their respective media, and 10 µL of the diluted cultures were dripped down the length of square TSA or TSAC Petri plates. After incubation at 37 °C overnight, colonies were counted to quantify bacterial concentrations.

### Quantification of antibiotic inhibitory concentrations

The inhibitory concentration for each antibiotic was determined by using a modified broth microdilution technique in 96-well plates, as described previously^36,37^. Briefly, *E. coli* and *E. faecalis* were grown in CDM supplemented with 1% glucose in the presence of serial dilutions of each antibiotic in triplicate. Bacterial OD_600nm_ was monitored with a plate reader (BioTek Synergy HTX) while the plates were incubated at 37 °C with continuous shaking. At a timepoint when untreated bacteria were in mid-to-late log phase (approximately 4 hours), the OD_600nm_ was recorded for all conditions, and these values were subjected to non-linear regression variable-slope analysis to determine the concentration of antibiotic that blocks 90% of bacterial growth (IC_90_).

### Statistical analysis and rigor of the data

All experiments were performed with at least three technical and three biological replicates. Statistical analysis and graphing were performed using GraphPad Prism (Version 10.4.1). Data are presented as means +/-standard deviation or standard error, as indicated and appropriate. Statistical significance was determined by using repeated measures two-way ANOVA followed by Bonferroni’s post hoc tests as indicated in the figure legends.

## RESULTS

### Oxygen quantification during logarithmic bacterial growth in planktonic cultures

Oxygen consumption is an indicator of respiration, which is a highly active pathway during logarithmic growth of aerobic bacteria^38,39^. To evaluate the capability of the Resipher to monitor oxygen consumption during active planktonic growth, we performed continuous measurements of dissolved oxygen in cultures of *Escherichia coli, Francisella novicida, Staphylococcus aureus*, and *Enterococcus faecalis* in rich media at 37 °C. Oxygen concentrations were monitored in 96-well plates inoculated with 10^6^ - 10^7^ CFU/mL using the Resipher, and, in parallel, growth of the bacteria was quantified from a second 96-well plate treated identically. All four species depleted oxygen from the media as bacterial growth progressed (Figure 1A-D). As expected, bacterial proliferation coincided with oxygen consumption, confirming that oxygen consumption serves as a robust proxy for bacterial growth. This result validates the ability of the Resipher to quantify oxygen in growing bacterial cultures.

**Figure 1.**
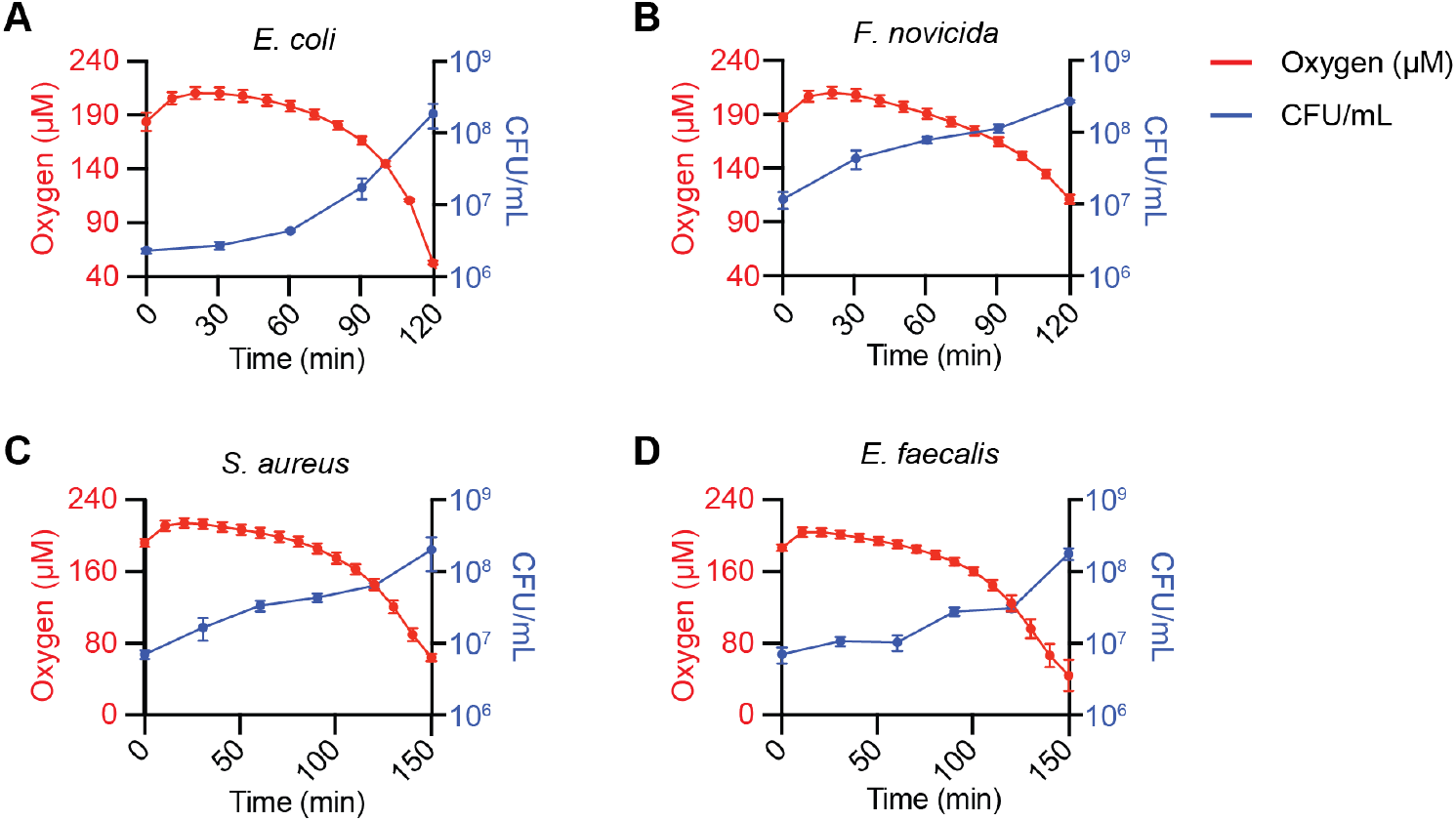
The Resipher quantifies oxygen concentration in the logarithmic phase of planktonic bacterial cultures. Time-dependent changes in oxygen concentration relative to bacterial colony-forming units in planktonic cultures of **(A)** *E. coli*, **(B)** *F. novicida*, **(C)** *S. aureus*, and **(D)** *E. faecalis*. Red lines indicate oxygen concentrations in cultures grown in TSB/TSBC at 37 °C and blue lines represent colony-forming units in a second plate simultaneously grown in identical conditions. Data shown are means +/-standard deviations in a single representative example of three biological replicates, each performed with N = 6 and 3 technical replicates for oxygen concentration and CFU enumeration, respectively.

### Establishing conditions to measure bacterial respiration independent of growth

Quantitative analysis of bacterial respiration can be confounded by concurrent changes in cell number, as oxygen consumption increases with proliferation. To accurately assess respiration as an intrinsic metabolic activity, it is necessary to establish conditions in which oxygen consumption can be measured independently of bacterial growth. To minimize proliferation while preserving metabolic activity, we reduced the incubation temperature from 37 °C to 25 °C and evaluated oxygen consumption across a range of bacterial densities. Lowering the temperature slows bacterial replication without abolishing respiratory activity, thereby enabling measurement of oxygen consumption over short time scales without significant changes in cell number.

In media lacking bacteria, the concentration of oxygen remained consistent at approximately 170 µM through the durations of the experiments, suggesting that the sensing probes themselves do not affect oxygen measurements. All four bacterial species displayed density-dependent consumption of oxygen from the culture media (Figure 2A-D). Among the tested species, *E. coli* and *F. novicida* depleted oxygen from the media more rapidly than the *S. aureus* and *E. faecalis*, and the densest cultures reached a steady-state plateau by 10 minutes. In all species, the most concentrated bacterial cultures (OD_600nm_ = 0.5 and 0.25) exhibited markedly lower oxygen concentration at the start of the measurements. This is likely due to oxygen depletion in the short time it takes to set up the experiment, suggesting that these densities of bacteria are unreliable for analysis. From these oxygen depletion profiles, we calculated OCR values that indicate greater oxygen consumption by *E. coli* and *F. novicida* than *S. aureus* and *E. faecalis* (Figure 2E-H). These results indicate that maximal depletion rates occur at or above OD_600nm_= 0.25, implying that this density may fall outside the linear range and therefore be suboptimal for analysis. At the lowest concentrations of *S. aureus* and *E. faecalis*, oxygen levels changed only minimally over the course of the experiments, indicating that their oxygen utilization was below the sensitivity threshold of the Resipher and may be influenced by the influx of ambient oxygen. Collectively, these observations indicate that intermediate oxygen consumption occurs at or just below OD_600nm_ = 0.25, supporting the use of this density for analysis at 25 °C.

**Figure 2.**
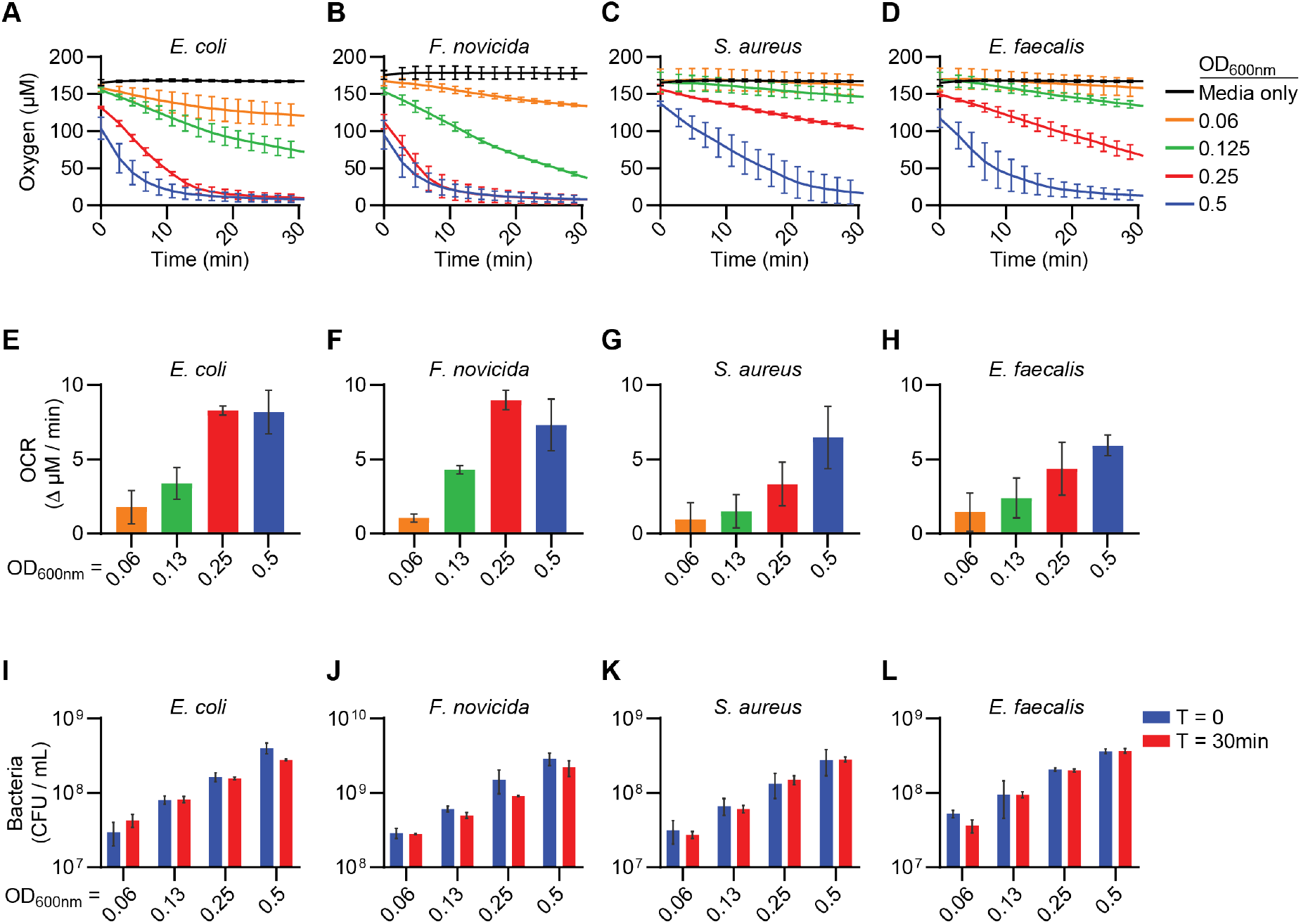
The Resipher measures oxygen consumption in conditions that limit changes in viable bacteria. Oxygen concentration measurements over time of **(A)** *E. coli*, **(B)** *F. novicida*, **(C)** *S. aureus*, and **(D)** *E. faecalis* incubated in TSB/TSBC at 25 ± 2 °C. **(E-H)** Initial oxygen consumption rates (OCR) of the four indicated species in the first 10 minutes of the experiments shown in (A-D). **(I-L)** Enumeration of colony forming units (CFU) before (T = 0) and after (T = 30 min) oxygen concentration measurements in (A-D) to evaluate the viability of the indicated species. Data presented are means +/-standard error means of three independent experiments, each with 6 technical replicates per condition.

To determine if the oxygen consumption profiles in these conditions were confounded by differences in bacterial concentration across the duration of these experiments, we quantified and compared the viable bacteria at the start and end of the oxygen measurement period by enumerating colony-forming units (CFU). CFU enumeration confirmed that bacterial numbers remained stable during the assay period, validating that measured oxygen consumption reflects metabolic activity rather than population expansion (Figure 2I-L). Taken together, these findings confirm the Resipher system as a sensitive platform for measuring oxygen consumption by bacteria under conditions where growth is minimized.

### Validation and reproducibility of high-throughput bacterial respiration measurements

To evaluate the accuracy and robustness of the Resipher for quantifying bacterial respiration, we compared oxygen consumption rates (OCR) in *E. coli* cultures obtained using the Resipher with those measured using the Oroboros O2k high-resolution respirometer, which employs a Clark-type polarographic oxygen sensor. Logarithmic-phase bacterial cultures were normalized to OD_600nm_ = 0.2 and analyzed in parallel using both platforms under comparable conditions. The average OCR across biological replicates was identical between the two methods (1.6 × 10^−6^ µM/min/CFU), with no statistically significant difference observed (Figure 3A). These findings demonstrate that the Resipher provides quantitatively equivalent measurements of bacterial oxygen consumption relative to a gold-standard respirometry approach.

**Figure 3.**
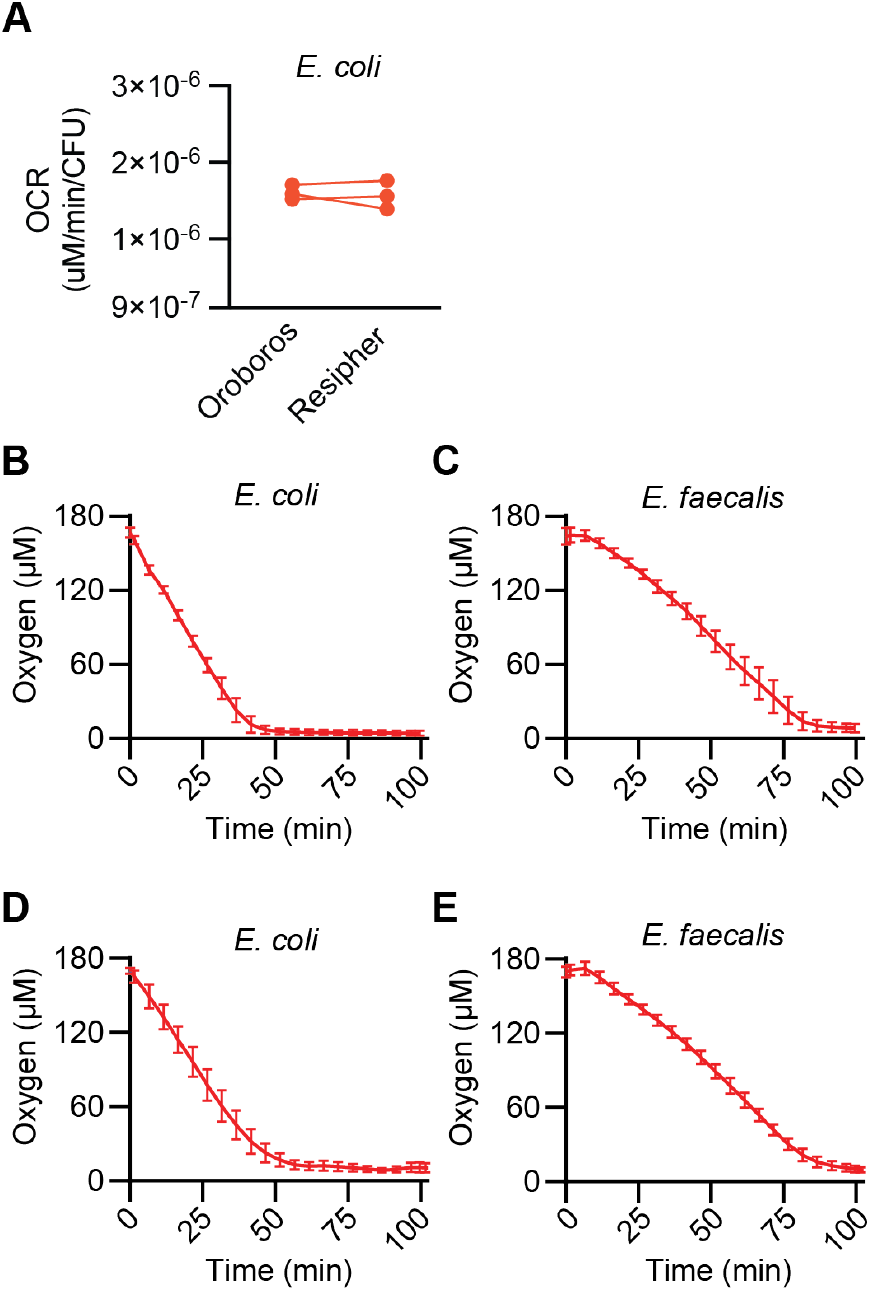
Validation and reproducibility of Resipher-based oxygen consumption measurements. **(A)** Comparison of oxygen consumption rates (OCR) measured in paired *E. coli* samples using the Resipher and the Oroboros O2k high-resolution respirometer. Each pair of data points represents measurements from the same biological sample analyzed on both platforms, with lines connecting paired measurements. No statistically significant differences were observed between the two methods. **(B, C)** Oxygen concentration measurements from a single biological replicate consisting of 24 technical replicates in cultures of **(B)** *E. coli* and **(C)** *E. faecalis* (OD_600nm_ = 0.2). Data represent means and standard deviations. **(D, E)** Oxygen concentration measurements from **(D)** *E. coli* and **(E)** *E. faecalis* (OD_600nm_ = 0.2) in three biological replicates performed on different days, each consisting of 24 technical replicates. Data represent means and standard errors.

To assess reproducibility, we evaluated both technical and biological variability using *E. coli* and *E. faecalis*. Oxygen concentrations recorded across 24 technical replicates per condition showed consistent oxygen consumption profiles, with standard deviations below 12 µM throughout the experiments (Figure 3B-3C). Independent biological replicates performed on separate days also demonstrated highly reproducible oxygen consumption dynamics, with standard errors below 13 µM (Figure 3D-3E).

Together, these results establish that the Resipher provides accurate, low-variability, and reproducible measurements of bacterial oxygen consumption, supporting its use as a reliable platform for high-throughput respiratory profiling.

### The Resipher detects nutrient-dependent changes in oxygen consumption

Bacteria rely solely on supplemented carbon sources in minimal media for metabolism and ATP production^8,40,41^. Thus, to determine if the Resipher can detect nutritionally induced changes in respiration, we supplemented minimal defined media with various carbon sources and measured the oxygen consumption in *E. coli* and *E. faecalis*. As shown previously, *E. coli* and *E. faecalis* display robust oxygen consumption in tryptic soy broth (Figure 4A and 4D). Studies on carbon source supplementation were performed in well-characterized minimal media for their respective bacterial species: M9 for *E. coli* and CDM for *E. faecalis*^34,35^. In minimal media lacking carbon sources, both *E. coli* and *E. faecalis* cultures exhibited negligible changes in oxygen over time (Figure 4A-D). This confirms that neither species can undergo aerobic respiration in these conditions due to an absence of substrates to fuel the ETC. The addition of carbon sources to minimal media enables oxygen consumption by both species. In *E. coli*, supplementing M9 minimal media with glucose or glycerol led to an initial burst of oxygen consumption in the first 25 minutes, after which oxygen utilization plateaus at approximately 140 µM (Figure 4A and 4C). In contrast, the *E. faecalis* oxygen depletion profiles demonstrated more prolonged oxygen consumption when CDM was supplemented with either carbon source (Figure 4B and 4D). This is potentially because CDM may contain the amino acids required for viability, thus eliminating the need for metabolically costly biosynthetic pathways. These results verify that the Resipher can monitor carbon source-induced changes in bacterial respiration.

**Figure 4.**
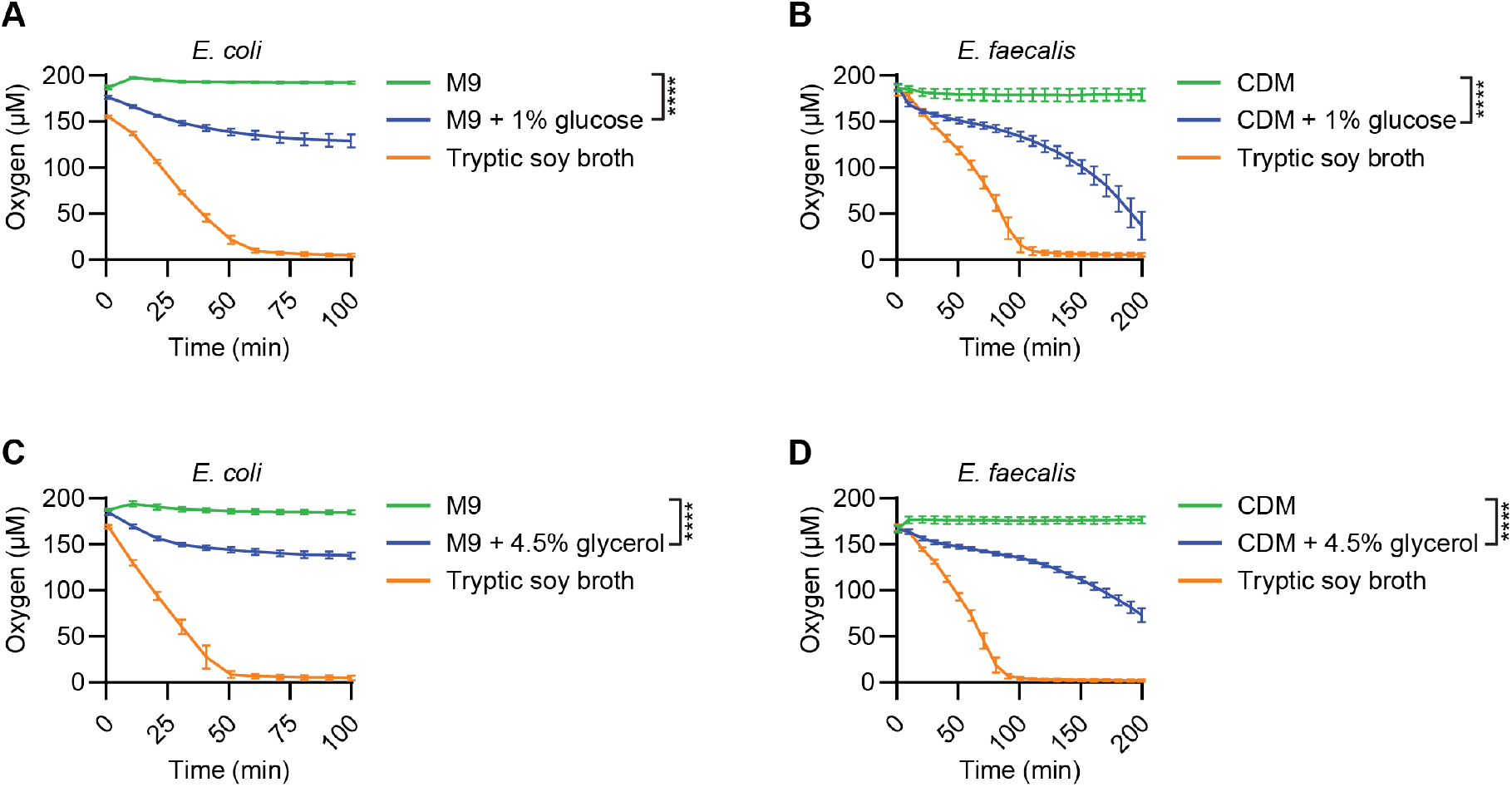
Supplementation of minimal-defined media with carbon sources is required for aerobic respiration. Oxygen concentration measurements of **(A, C)** *E. coli* and **(B, D)** *E. faecalis* in rich media (tryptic soy broth) or the indicated minimal defined media (M9 or CDM) supplemented with **(A, B)** glucose or **(C, D)** glycerol. Data shown are a single representative example (N = 6 technical replicates) of three biological replicates. Two-way ANOVA statistical tests with Bonferroni’s multiple comparisons were performed to assess differences between treated and untreated conditions. (**** P < 0.0001)

### Electron transport chain inhibitors modulate oxygen consumption

Inhibitors of various components of the ETC elicit distinct responses in respiratory activity. Although the effects of these inhibitors on mitochondrial oxidative phosphorylation are well-characterized, their effects on bacterial respiration have received less attention^42^. We employed the Resipher to assess the impact of ETC inhibitors on *E. coli* and *E. faecalis* oxygen consumption in a high-throughput format. As expected, treatment of both *E. coli* and *E. faecalis* with 10 mM CCCP increased oxygen consumption (Figure 5A and 5B)^43,44^. Consistent with previous reports, treatment with higher concentration of CCCP (100 mM) led to blockage of oxygen consumption, likely due to generation of reactive oxygen species and cell death (data not shown)^45,46^.

**Figure 5.**
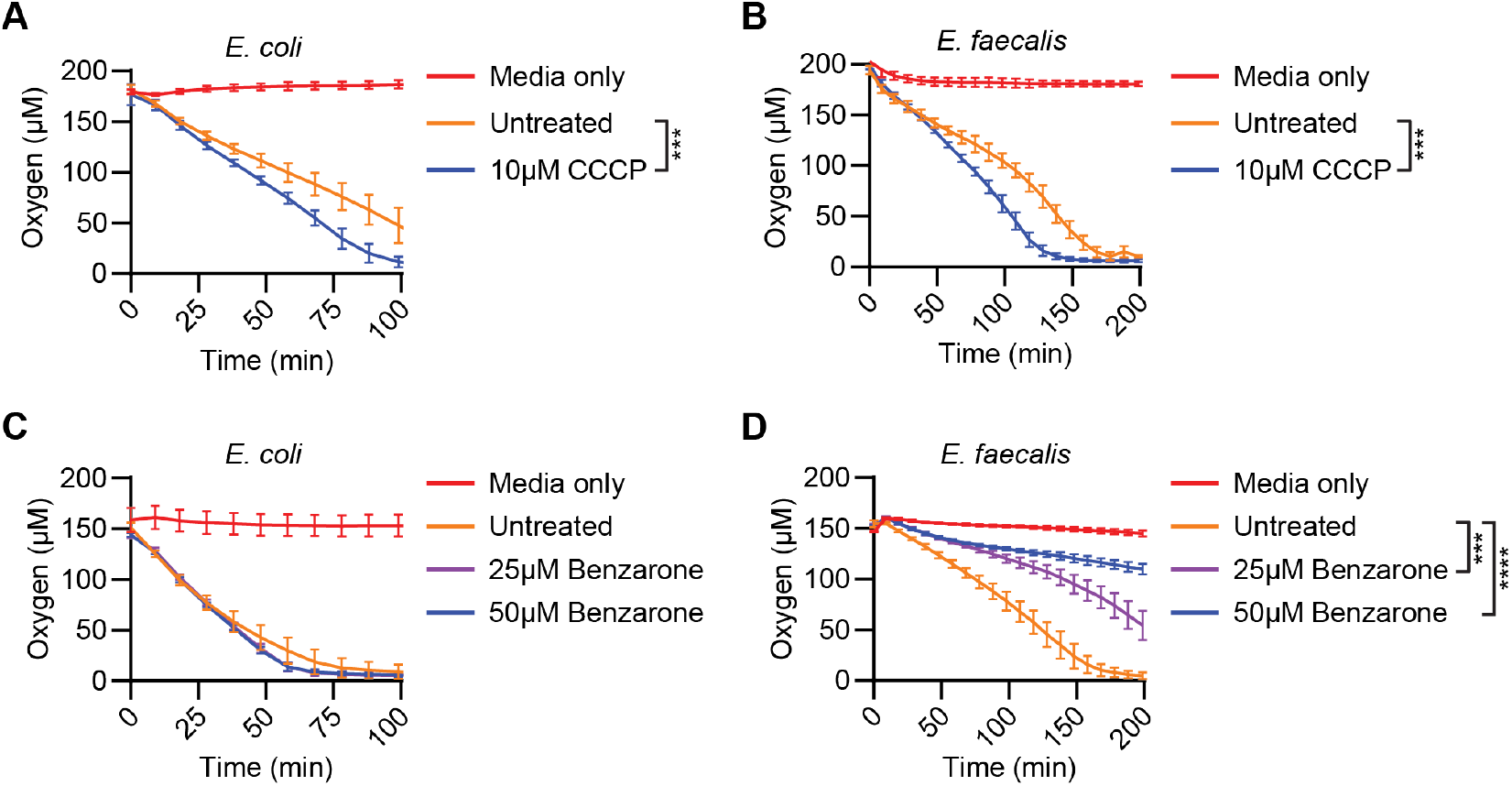
Electron transport chain inhibitors have distinct effects on oxygen consumption by *E. coli* and *E. faecalis*. Oxygen consumption measurements of **(A, C)** *E. coli* and **(B, D)** *E. faecalis* in chemically defined media (CDM) + 1% glucose following treatment with **(A, B)** CCCP and **(C, D)** benzarone at the indicated concentrations. Data shown are a single representative example of three biological replicates and are composed of 6 technical replicates each. Two-way ANOVA statistical tests with Bonferroni’s multiple comparisons were performed to assess differences between treated and untreated conditions. (*** P = 0.0002, **** P < 0.0001)

Benzbromarone and its active metabolite, benzarone, potently inhibit mitochondrial oxidative phosphorylation; however, relatively little is known about their effects on bacterial respiration^47,48^. Given this paucity of data, we employed the Resipher to determine how benzarone affects *E. coli* and *E. faecalis*. Neither 25 µM nor 50 µM benzarone affected *E. coli* oxygen consumption (Figure 5C). In contrast, benzarone had a dose-dependent effect on *E. faecalis* (Figure 5D). 25 mM benzarone modestly attenuated oxygen consumption approximately 50% and 50 mM blocked up to 70%, following 150 minutes of exposure. These data are consistent with a recent report demonstrating benzbromarone inhibits growth of Gram-positive bacteria but not Gram-negative organisms^49^. Based on these results, the antibacterial activities of benzarone and benzbromarone on Gram-positive bacteria are likely through blocking bacterial respiration.

### Bactericidal and bacteriostatic antimicrobials cause distinct changes in respiration

Distinct respiratory responses are associated with bactericidal and bacteriostatic antibiotics. In some conditions, bactericidal antibiotics induce a transient increase in oxygen consumption, and, in contrast, bacteriostatic antibiotics generally suppress metabolic activity and lead to attenuated oxygen consumption^11,12,20^. Thus, we employed the Resipher to determine if it is capable of detecting these distinct profiles. Inhibitory concentrations of the antibiotics were determined by generating dose-response curves and analyzing the data with non-linear regression (Figure S1). We exposed *E*.*coli* and *E. faecalis* to enrofloxacin and ampicillin to quantify bactericidal antibiotic-induced changes in oxygen consumption. Significant inhibition of oxygen consumption was observed in *E. coli* and *E. faecalis* following treatment with concentrations of enrofloxacin 5-10-fold above the IC_90_ (Figures 6A and 6B). A sub-inhibitory concentration of enrofloxacin caused a modest but statistically significant increase in respiratory activity in *E. faecalis*, whereas *E. coli* oxygen consumption remained unaffected under the same condition. Similarly, sub-inhibitory concentrations of ampicillin (1/8 IC_90_) resulted in elevated respiratory activity in both *E. coli* and *E. faecalis* (Figures 6C and 6D). These results are consistent with prior observations suggesting activation of compensatory metabolic pathways in response to antibiotic-induced bacterial stress_12,20,50_.

**Figure 6.**
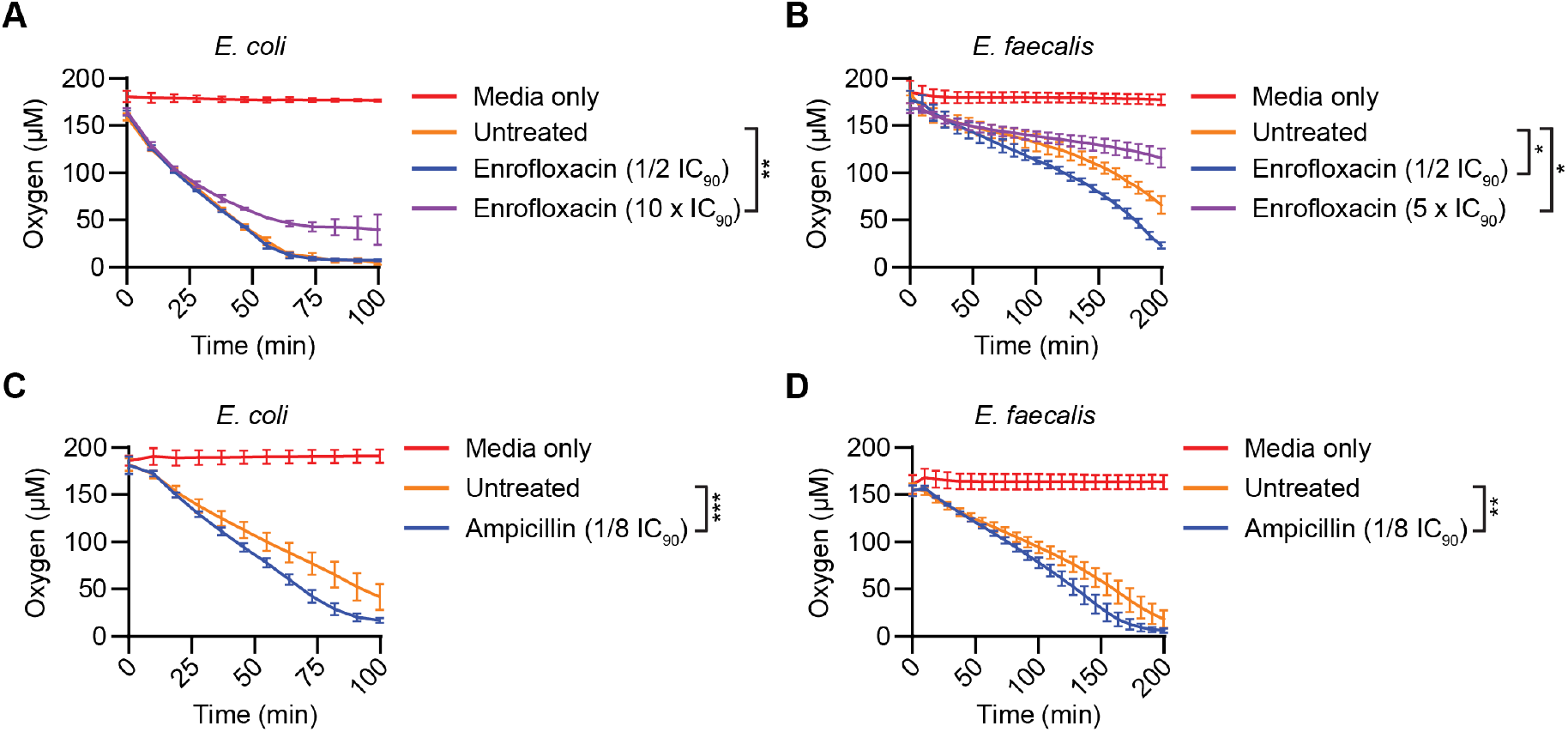
Treatment with subinhibitory concentrations of bactericidal antibiotics enhances oxygen consumption. Treatment of **(A, C)** *E. coli* and **(B, D)** *E. faecalis* with **(A, B)** enrofloxacin and **(C, D)** ampicillin at concentrations relative to their calculated IC_90_ in CDM + 1% glucose. Statistical comparisons between treated and untreated groups were performed using two-way ANOVA and Bonferroni’s multiple comparison tests. Data shown are a single representative example of at least three biological replicates and are composed of 6 technical replicates each. (* P < 0.05, ** P < 0.01,*** P < 0.001)

Unlike bactericidal antimicrobials, bacteriostatic drugs are known to lead to decreased metabolism and oxygen consumption^12,20^. To determine if the Resipher can detect these effects, we measured oxygen consumption following treatment with inhibitory concentrations of doxycycline, chloramphenicol, and tetracycline. Exposure to these bacteriostatic antimicrobials caused a marked reduction in oxygen consumption by both *E. coli* and *E. faecalis* to various degrees (Figures 7A-7F). Contrary to our results with bactericidal antimicrobials, testing bacteriostatic antibiotics at subinhibitory concentrations diminished oxygen consumption in a dose dependent fashion (Figure S2). Taken together, these results support the utility of oxygen consumption as an indicator of metabolic activity and validate the use of the Resipher in measuring oxygen in bacterial cultures.

**Figure 7.**
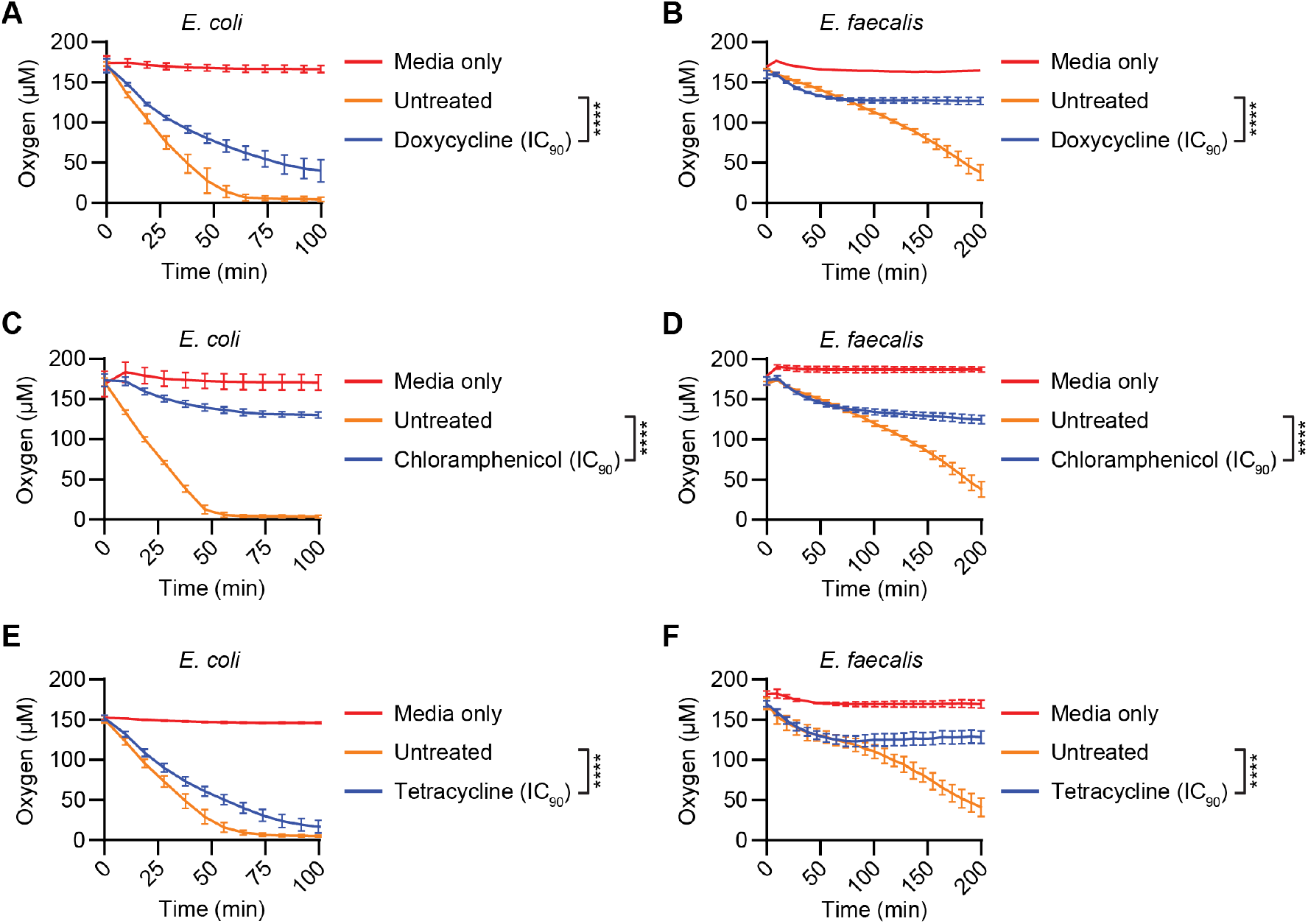
Bacteriostatic antibiotics attenuate oxygen consumption by *E. coli* and *E. faecalis*. Treatment of *E. coli* and *E. faecalis* with the bacteriostatic antibiotics **(A, B)** doxycycline, **(C, D)** chloramphenicol and **(E, F)** tetracycline at their IC_90_s in CDM + 1% glucose. Statistical comparisons between treated and untreated groups were performed using two-way ANOVA model with Bonferroni’s multiple comparison tests. Data shown are a single representative example of greater than three biological replicates and are composed of 6 technical replicates each. (**** P < 0.0001)

## DISCUSSION

Quantitative measurement of bacterial respiration provides a functional window into cellular metabolic state, stress adaptation, and antimicrobial response. In this study, we establish a high-throughput approach for real-time profiling of bacterial oxygen consumption by adapting the Resipher platform for microbial systems. Importantly, oxygen consumption rates measured using the Resipher were quantitatively comparable to those obtained with a Clark-type electrode-based high-resolution respirometry system, supporting the accuracy of this platform relative to established methodologies. Beyond technical validation, our findings demonstrate that oxygen consumption serves as a sensitive and dynamic phenotypic readout that captures biologically meaningful responses to nutrient availability, electron transport chain perturbation, and antibiotic exposure across diverse bacterial species. These results position respiratory profiling as a broadly applicable strategy for interrogating bacterial physiology.

In exponential-phase planktonic cultures, oxygen consumption closely reflected the bacterial growth. Previous studies have similarly shown that oxygen depletion correlates with biomass accumulation in bacterial species, highlighting the reliability of oxygen consumption as an indicator of bacterial growth^51,52^. Unlike traditional endpoint assays, the Resipher allows continuous monitoring of oxygen consumption, providing real-time insights into respiratory dynamics that can be correlated with bacterial viability during logarithmic-phase growth.

At a suboptimal temperature that limits bacterial growth (25 °C), the Resipher reliably measures oxygen consumption. This capability is essential for respiration assays, where microbial growth and proliferation can confound metabolic measurements^53,54^. Although, reduced temperature can influence multiple physiological processes, including enzyme activity, membrane properties, and respiratory flux, this was necessary to make certain that oxygen consumption rates were in the linear and interpretable range while eliminating the confounding effect of bacterial growth during the study. Because of this, care should be taken when interpreting physiological properties that are known to be temperature sensitive, as the respiratory activity measured at 25 °C may not fully reflect optimal growth conditions. By optimizing the bacterial density, we identified conditions where oxygen concentration was readily measurable. These data establish OD_600nm_ below 0.25 as an acceptable range for quantitative assessments of bacterial oxygen consumption using this system. However, experiments can be performed at lower optical densities to increase the duration of the assays and allow for slower oxygen depletion.

Our optimization studies also revealed a practical limitation of respiratory profiling at very high bacterial densities. Cultures approaching stationary-phase densities rapidly depleted dissolved oxygen, often reaching minimal oxygen concentrations before stable measurements could be obtained. As a result, these conditions fall outside the optimal dynamic range for quantitative oxygen consumption analysis. This phenomenon is not unique to the Resipher and represents a general limitation of oxygen consumption measurements, as rapid oxygen depletion can compromise the ability of any respirometry platform to accurately resolve respiratory differences in highly concentrated samples. Accordingly, respiratory profiling is most informative when performed at bacterial densities that permit gradual oxygen consumption throughout the measurement period.

Nutrient availability and substrate specificity are key determinants of bacterial respiration. Our results highlight the strict dependence of *E. coli* and *E. faecalis* on exogenous carbon sources for aerobic metabolism in minimal media. Oxygen consumption was negligible in both species when carbon sources were absent, in agreement with prior studies demonstrating that ETC activity requires metabolic fuel^16,55^. The restoration of respiration following carbon supplementation underscores the essential role of nutrient availability in driving bacterial energy metabolism.

Bacteria exhibit distinct respiratory responses to different carbon sources, and these data suggest that the Resipher is sufficiently sensitive to detect nutrient-dependent changes in respiration under controlled conditions. The intricate interplay between nutrient availability, metabolic prioritization, and respiratory regulation can significantly influence bacterial respiration, and the Resipher can provide valuable insight into these physiological responses.

The Resipher also enabled profiling of respiration in response to ETC inhibitors. The ETC establishes a proton gradient across the bacterial cytoplasmic membrane and the resulting electrochemical gradient drives ATP generation through the activity of ATP synthase. ETC-driven oxygen consumption is restricted by proton translocation through ATP synthase, but protonophores, such as carbonyl cyanide m-chlorophenylhydrazone (CCCP), facilitate proton leakage and accelerate both proton flow and reduction of oxygen to water^43,44^. Consistent with our expectations, CCCP treatment increased oxygen consumption in both *E. coli* and *E. faecalis*, reflecting an uncoupled respiratory state characterized by increased oxygen consumption^44^. In contrast, benzarone selectively inhibited oxygen consumption in *E. faecalis* but not in *E. coli*, suggesting potential Gram-positive-specific activity. These data align with prior reports showing that benzbromarone specifically affects Gram-positive but not Gram-negative organisms^49^. Thus, these findings suggest that oxygen consumption profiling could be useful for identifying compounds that alter bacterial respiration, although broader validation across additional inhibitor classes will be required^56^.

This work also demonstrates that how the Resipher distinguishes between bactericidal and bacteriostatic antibiotics based on their distinct effects on oxygen consumption. Bactericidal antibiotics such as enrofloxacin and ampicillin transiently increased oxygen consumption at sub-inhibitory concentrations, likely reflecting activation of stress-response pathways and compensatory metabolism, as reported previously^12,20^. In contrast, bacteriostatic antibiotics including chloramphenicol and tetracycline suppressed oxygen consumption, consistent with their known inhibitory effects on protein synthesis and cellular metabolism. These divergent profiles validate the distinct metabolic consequences of bactericidal and bacteriostatic antibiotics and support the use of OCR as a functional physiological readout associated with antibiotic activity. These findings support oxygen consumption as a functional physiological phenotype associated with antibiotic-induced metabolic responses that may complement traditional growth-based antimicrobial susceptibility assays.

Care must be placed in interpreting the results of oxygen consumption measurements when using inhibitors, as the drugs themselves may affect dissolved oxygen in the media, irrespective of the presence of bacteria. Some drugs, such as pyrogallol, spontaneously generate superoxide and affect dissolved oxygen^57,58^. Conversely, decomposition of hydrogen peroxide and other peroxides leads to aberrant measurements that are unrelated to bacterial respiration^59^. The Resipher can readily discriminate between these chemical artifacts and biologically relevant changes. Since the Resipher can perform measurements in a high-throughput fashion, it has the capacity for parallel, multi-well measurements to facilitate rigorous assessments, helping to separate compound-specific chemical reactivity from genuine metabolic responses.

In summary, this work suggests that high-throughput respiratory profiling can provide biologically informative measurements of bacterial physiology across diverse species and experimental conditions. By linking oxygen consumption to nutrient utilization, electron transport chain activity, and antibiotic responses, our findings support respiration as a versatile functional phenotype that complements traditional growth-based analyses. While the Resipher provides the technical framework for these measurements, the broader significance of this work lies in enabling scalable investigation of bacterial bioenergetics, stress adaptation, and antimicrobial action. As interest in microbial metabolism continues to expand, respiratory profiling has the potential to become a valuable approach for studying bacterial physiology in clinical, environmental, and engineered systems.

## AUTHOR CONTRIBUTIONS

NK: Validation and investigation. NK and AE: Methodology, formal analysis, data curation, writing – original draft, writing – reviewing & editing, visualization, and project administration. AE: Conceptualization, resources, supervision, and funding acquisition.

## ACKNOWLEDGEMENTS

We wish to thank Walker Inman, Richard Bryan, Kin Lo, Silke Grainger, and Ryan Titmas (Lucid Scientific) for providing technical knowledge and access to the Resipher platform. Additionally, we acknowledge Chloe Van Horn and Leah Eshraghi for critically reviewing the manuscript and Adriana G Morales Rivera, Jose A Lemos, Feng Yue and Junxiao Ren for providing access to technical resources, assistance, and training.

## SUPPLEMENTARY FIGURES AND TABLE

**Figure S1.**
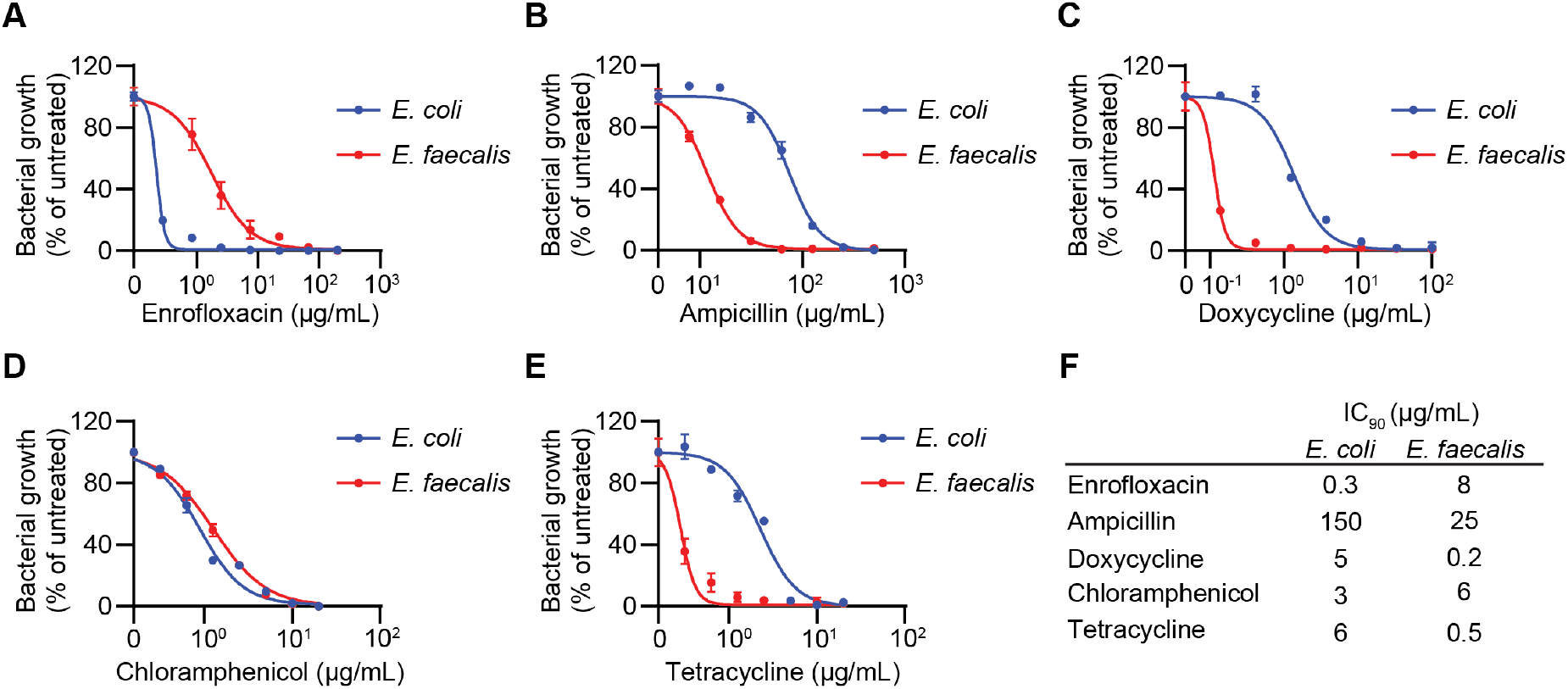
Sensitivity of *E. coli* and *E. faecalis* to antibiotics. Growth of *E. coli* and *E. faecalis* in CDM + 1% glucose after treatment with **(A)** enrofloxacin, **(B)** ampicillin, **(C)** doxycycline, **(D)** chloramphenicol and **(E)** tetracycline. Non-linear regression curves using a four-parameter variable-slope model were plotted to determine the inhibitory concentrations of each antibiotic against both bacterial strains. Data are representative of three biological replicates, each containing three technical replicates. **(F)** Calculated IC_90_ concentrations of the antibiotics in (A-E) for *E. coli and E. faecalis*.

**Figure S2.**
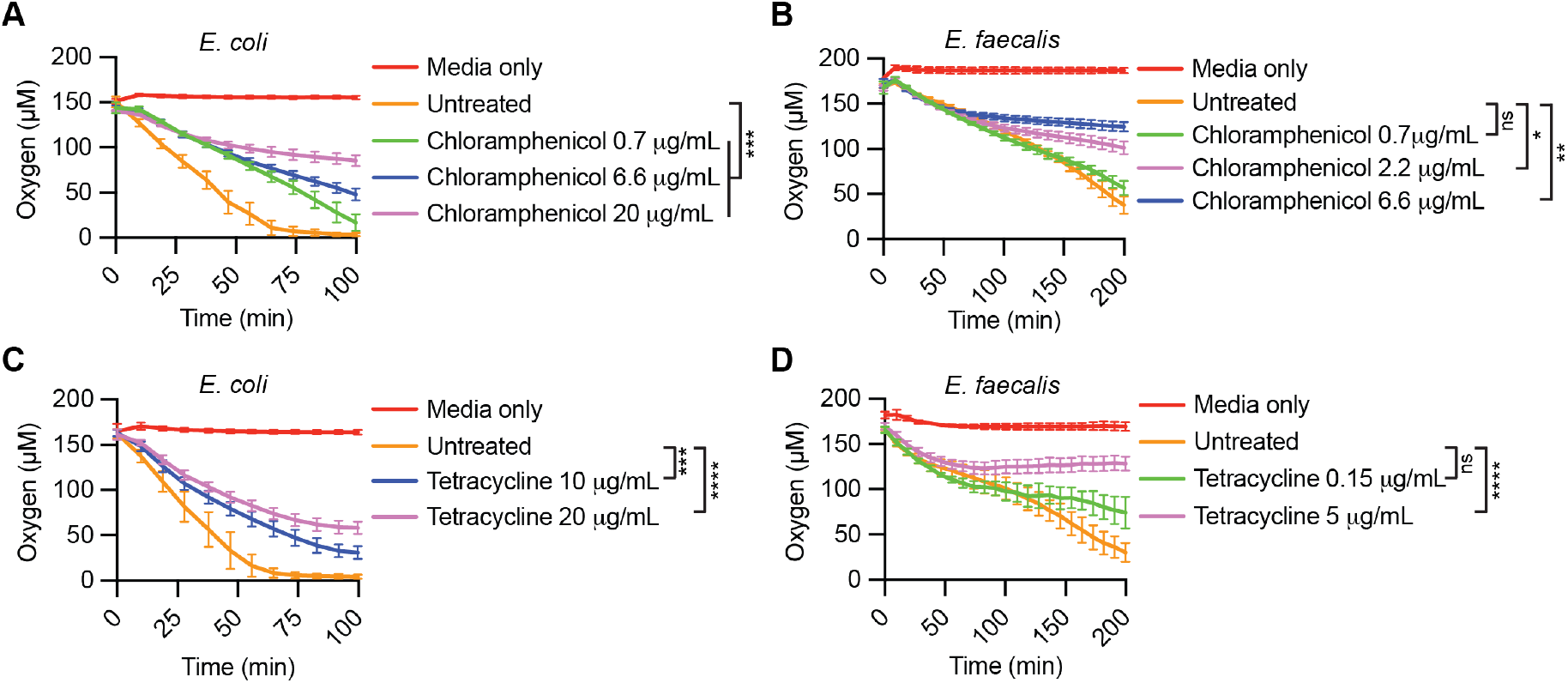
Dose-dependent changes in oxygen consumption in response to bacteriostatic antibiotics. Oxygen consumption measurements of **(A, C)** *E. coli* and **(B, D)** *E. faecalis* in chemically defined media (CDM) + 1% glucose following treatment with **(A, B)** chloramphenicol and **(C, D)** tetracycline at the indicated concentrations. Data shown are a single representative example of three biological replicates and are composed of 6 technical replicates each. Two-way ANOVA statistical tests with Bonferroni’s multiple comparisons were performed to assess differences between treated and untreated conditions. (* P < 0.05, ** P < 0.01, *** P < 0.001, **** P < 0.0001)

**Table S1.**
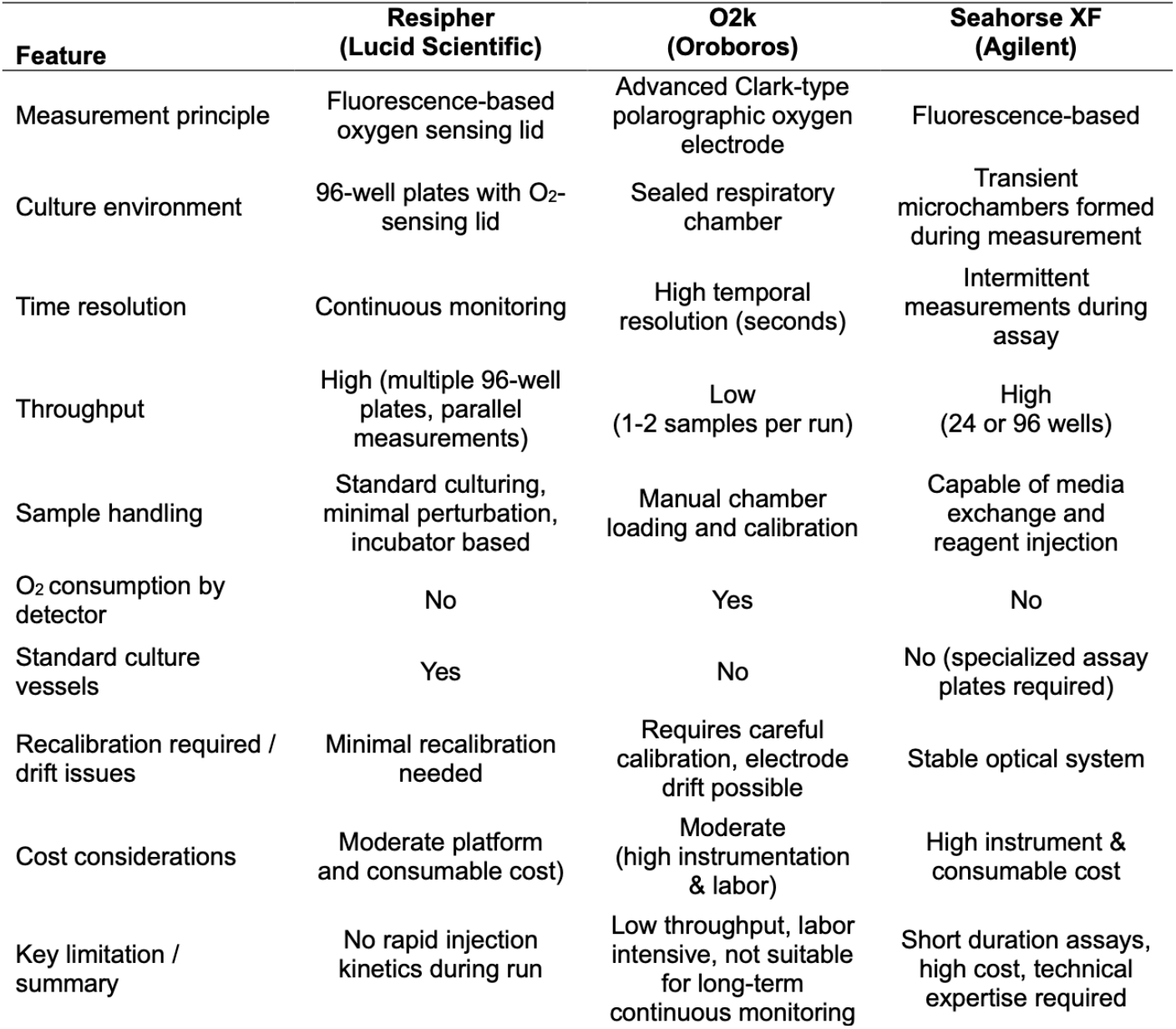
Comparison of platforms commonly used to measure oxygen consumption.

| **Feature** | **Resipher**<br>**(Lucid Scientific)** | **O2k**<br>**(Oroboros)** | **Seahorse XF**<br>**(Agilent)** |
| --- | --- | --- | --- |
| Measurement principle | Fluorescence-based oxygen sensing lid | Advanced Clark-type polarographic oxygen electrode | Fluorescence-based |
| Culture environment | 96-well plates with O<sub>2</sub>-sensing lid | Sealed respiratory chamber | Transient microchambers formed during measurement |
| Time resolution | Continuous monitoring | High temporal resolution (seconds) | Intermittent measurements during assay |
| Throughput | High (multiple 96-well plates, parallel measurements) | Low (1-2 samples per run) | High (24 or 96 wells) |
| Sample handling | Standard culturing, minimal perturbation, incubator based | Manual chamber loading and calibration | Capable of media exchange and reagent injection |
| O<sub>2</sub> consumption by detector | No | Yes | No |
| Standard culture vessels | Yes | No | No (specialized assay plates required) |
| Recalibration required / drift issues | Minimal recalibration needed | Requires careful calibration, electrode drift possible | Stable optical system |
| Cost considerations | Moderate platform and consumable cost | Moderate (high instrumentation & labor) | High instrument & consumable cost |
| Key limitation / summary | No rapid injection kinetics during run | Low throughput, labor intensive, not suitable for long-term continuous monitoring | Short duration assays, high cost, technical expertise required |

